# Bioactive Natural Product Discovery via Deuterium Adduct Bioactivity Screening

**DOI:** 10.1101/2023.03.16.532988

**Authors:** N.A. Zill, Y. Du, S. Marinkovich, D. Gu, J. Seidel, W. Zhang

## Abstract

The discovery of bioactive natural products lies at the forefront of human medicine. The continued discovery of these molecules is imperative in the fight against infection and disease. While natural products have historically dominated the drug market, discovery in recent years has slowed significantly, partly due to limitations in current discovery methodologies. This work demonstrates a new workflow, Deuterium Adduct Bioactivity Screening (DABS), which pairs untargeted isotope labeling with whole cell binding assays for bioactive natural product discovery. DABS was validated and led to the discovery of a new isoprenyl guanidine alkaloid, zillamycin, which showed anti-cancer and anti-microbial activities. DABS thus represents a new workflow to accelerate discovery of natural products with a wide range of bioactive potential.

## Introduction

Natural products (NPs) have a long and impressive history as sources of pharmaceutical compounds and as important biological research tools. From 1981 to 2019, 65.4% of the new FDA approved drugs are NPs, their mimics, or their derivatives^1^. Particularly compared to combinatorial compounds, analysis of physicochemical properties highlights that NPs occupy a much larger and distinct area in molecular space, and are evolutionarily optimized as drug-like molecules due to their intrinsic ability to interact with biological targets^2,3,4^.

Despite the past success and continuing interest in NP-based drug discovery, modern NP research has witnessed slow progress in discovery of promising drug leads, partly due to limitations in current discovery methodologies^5^. Historically, the most widespread discovery process is based on phenotypic assays, particularly cell growth inhibitory assays, which has several notable drawbacks such as low sensitivity and flexibility^6^. Alternatively, microbial genome mining offers a modern method to discover NPs with potentially new chemical structures, but their bioactivities are often elusive^7,8^. Considering that new technologies, such as metabolomics^9,10,11,12^ and synthetic biology^13,14^ have shortened the time for NP identification and production^15^, there is an urgent need to develop new screening assays for bioactive NP discovery from complex mixtures.

We here employ whole cell binding assays for primary bioactive NP screening, a method expected to have higher sensitivity, throughput, and flexibility than the traditional phenotype-base screening methods. More importantly, these reporting cells simultaneously enrich active components from pre-binding NP mixtures as shown by the intracellular quantification studies of bioactive compounds by us and other researchers^16,17,18^. These assays thus allow facile active compound re-isolation from cells and detection by sensitive metabolomics analysis. Doing so, any metabolite that has a specific target in reporting cells may be enriched and revealed, significantly enhancing the rate of active compound discovery. This method is expected to be less sensitive to the amount of active compound in the pre-binding mixtures, and thus is particularly efficient in identifying compounds that are missed during the traditional phenotypic assay-based screening.

To facilitate the mass spectrometry (MS)-based identification and compound tracking in whole cell binding assays, untargeted isotope labeling of the NP mixture is preferred^19,20,21,22^. One useful tag is the C-D bond, which is expected to minimally affect the chemical and biological properties of labeled NPs. Deuterium may be readily incorporated into NPs through natural metabolic processes by growing NP producing organisms in D_2_O^23^. Previous work has illustrated that microbial organisms may tolerate high concentrations of D_2_O^24^, suggesting feasible culturing conditions to label a broad spectrum of metabolites in a scaffold-independent manner.

In the present study, we developed and implemented a new Deuterium Adduct Bioactivity Screening (DABS) workflow (**Figure 1**) combining untargeted isotope labeling and whole cell binding assays for bioactive NP discovery. DABS was validated using known bioactive NPs, and further led to the discovery of a new cytotoxic natural product from a well-studied organism of *Streptomyces griseofuscus*.

**Figure 1:**
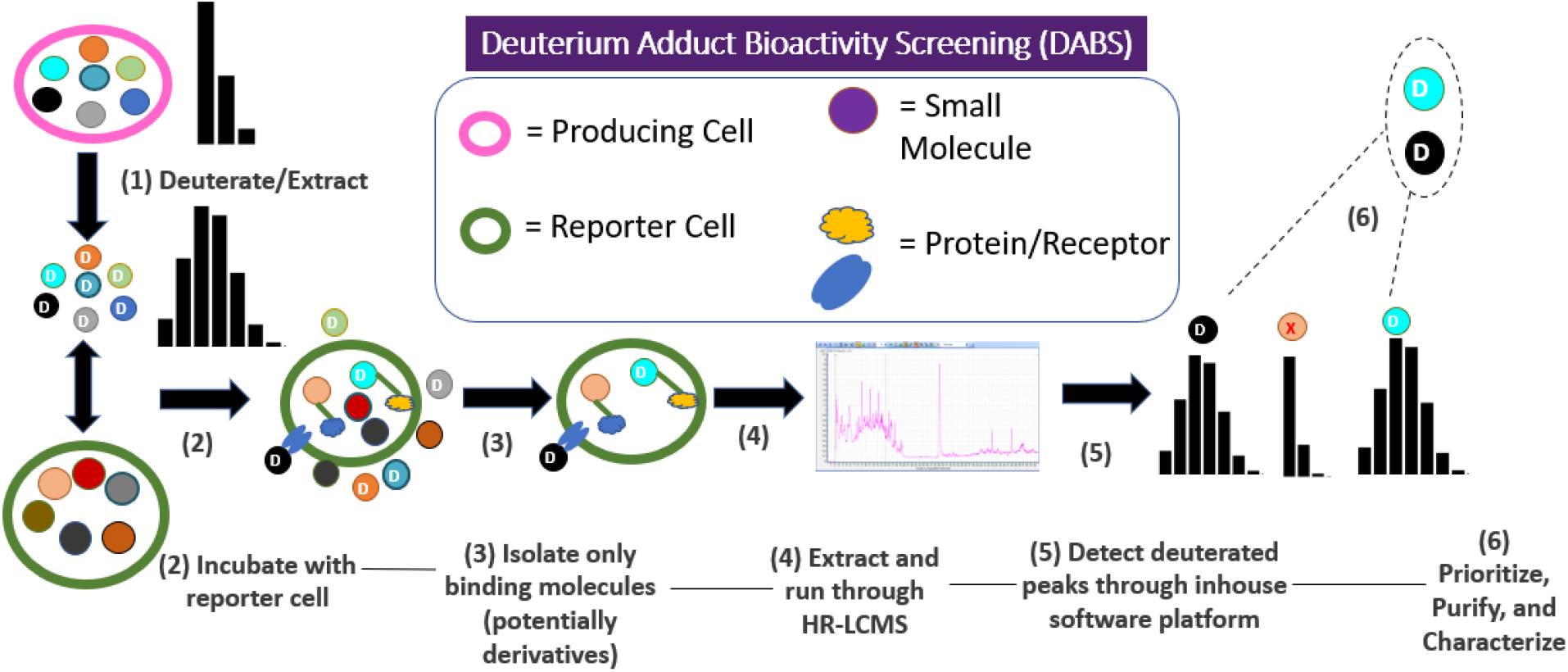
Deuterium Adduct Bioactivity Screening workflow. NP producing microbial strains are grown in deuterated water in order to label the metabolome. These metabolites are extracted and then incubated in a cell binding assay with a cell line of choice (microbial or mammalian). The cell line is washed to remove non-binders and the metabolome is re-extracted. Metabolites originating from the deuterated extract will be readily detectable due to their unique deuterated isotopic signature.

## Results and Discussion

### Untargeted Deuterium Labeling of NPs

To better understand the sparsely studied effect of D_2_O on microbial metabolomes and its viability as a labeling strategy, we selected a model actinomycete, *Streptomyces albus* J1074, to observe its growth and metabolic profile in D_2_O. This strain has a well-studied growth condition and rich secondary metabolome^25,26,^, allowing for a facile examination of deuterium toxicity and metabolic shifts. In terms of burden to the cell, modest growth lag effects were observed when cultures were grown on Mannitol-Soya (MS) agar plates at various D_2_O concentrations (0, 25%, 50%, 75%). Starting with a native full lawn sporulation time of ∼ one week, the D_2_O added approximately 1-2 extra days of growth per 25% addition of D_2_O before observing the full lawn sporulation. Inoculation cultures from deuterated plates tended to reduce the time delay in further growth experiments. No other variances, such as color and final cell density, were noted. We next probed the small-molecule metabolome by Liquid Chromatography – High Resolution Tandem Mass Spectrometry (LC)-HRMS-MS_n_ analysis, and observed that the total ion chromatograms (TICs) upon reaching sporulation varied more significantly with increasing D_2_O concentrations, while a minimal impact was observed up to 25% of D_2_O (**Figure S1**). To assess metabolomic incorporation of deuterium, we zoomed in on a family of well-known NPs of *S. albus*, the depsipeptide antimycins, and observed an increased deuterium incorporation with increasing concentrations of D_2_O (**Figure 2a**).

**Figure 2:**
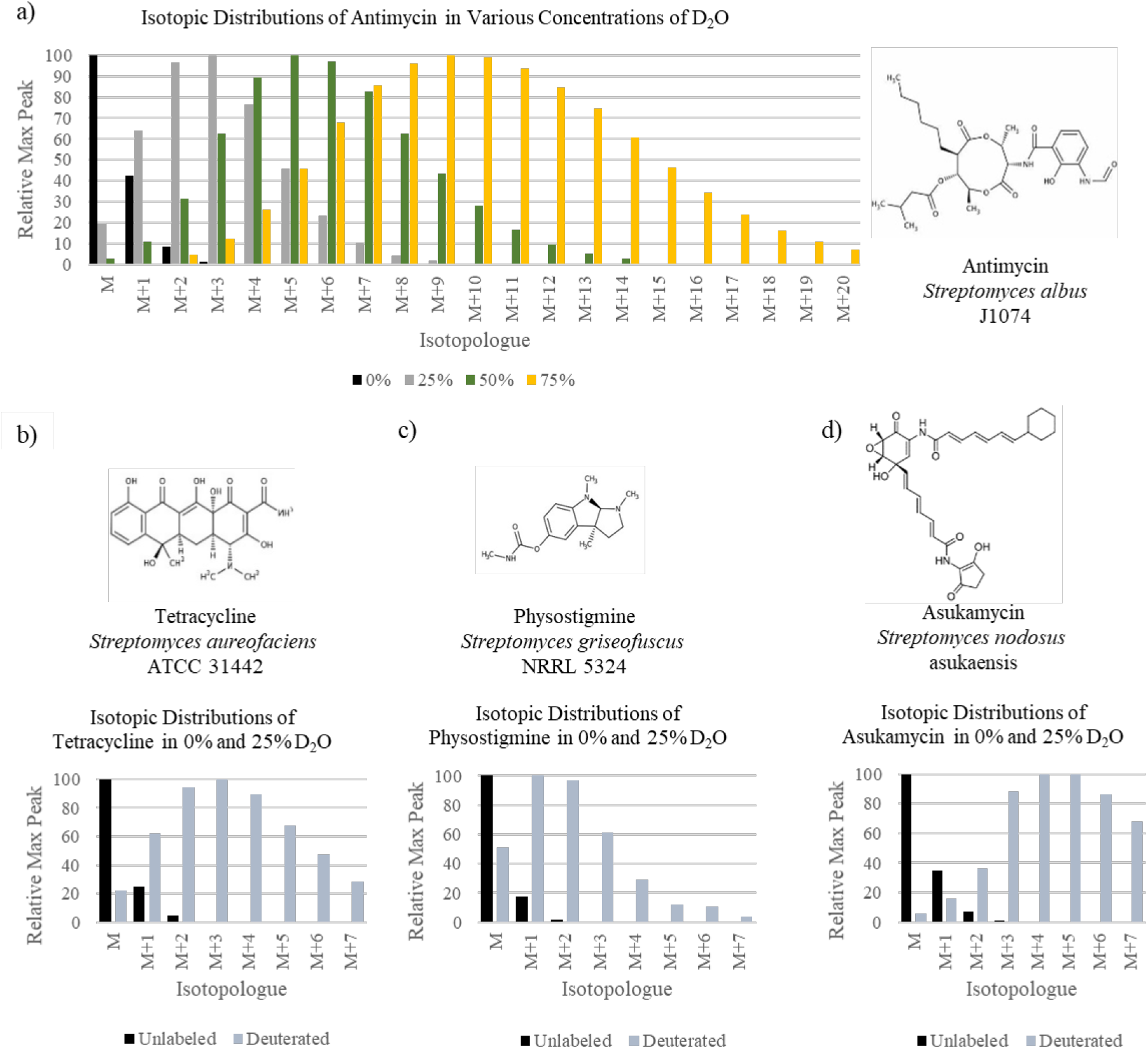
Deuterium’s effect on metabolomes. A varirety of *Streptomyces* strains were grown in various concentrations of D_20_. The agar plate extracts were screened through LC-MS showing a) a variety of D_2_O concentrations with Antiniycin (predicted m/z=549.28, observed m/z=549.27), and the isotopologue variations produced by 25% D_2_O of b) tetracycline (predicted m/z=445.16n, observed m/z=445.16), c) physostigmine (predicted m/z=276.17, observed m/z=267.17), and d) asukainycin (predicted m/z=547.24, observed m/z=547.24). “M” represents the in/z peak corresponding to the unlabeled base peak.

For automated detection of deuterated metabolites, a software program dubbed “D_2_O Sorter” was written as detailed in the methods section. In short, “D_2_O Sorter” was generated using a list of molecular formulas expected to be physically possible given constraints such as the Lewis rules and electron configurations. From these formulas, the expected isotope distribution of both undeuterated and deuterated metabolites was generated. Comparing the two spectra, an evident trend was detected that allowed sorting of the deuterated metabolites by comparing the relative (M+1) and (M+2) values of each metabolite relative to the base (M) peak within a given range of m/z. This allowed for highly fidelitous sorting of the deuterated metabolites from complex mixtures. “D_2_O Sorter” was applied to the culture extracts of *S. albus* and as expected, deuterated antimycins were readily detected by the program.

Based on the promising results of untargeted deuterium labeling of NPs in *S. albus*, we next tested additional *Streptomyces* strains with various known NPs for further validation. We selected a D_2_O concentration of 25% to minimize the growth inhibition and metabolic shift of producing organisms while labeling the majority of produced NPs. The three tested *Streptomyces* strains showed the similar overall trends in both growth and metabolic profile shift in the presence of D_2_O as *S. albus*. The selected known NPs from these strains showed clear incorporation of deuterium into their scaffolds at 25% D_2_O (**Figure 2b-d**). Notably, the deuterated metabolites showed slight drift in retention time on a C18 column, more apparently with higher mass isotopologues, which is consistent with literature illustrating the increased polarity of C-D vs C-H bonds^27^. Each strain had well over 50 small molecules predicted to be deuterated, demonstrating a wealth of metabolites deuterated in these rich producers of natural products.

### Deuterium Adduct Bioactivity Screening (DABS) Workflow and Validation

In a typical DABS workflow as shown in **Figure 1**, an NP-producing strain is grown in 25% D_2_O and extracted to generate a complex mixture of labeled natural products. This extract is applied to a reporter cell, such as microbial or mammalian cells, and non-specific or weak-binding compounds will be washed away. The strong-binding compounds will be locally concentrated in live cells and re-isolated from these cells through chemical extraction followed by LC-HRMS- MS^n^ detection and computational data analysis. Specifically, the isotopic feature readily distinguishes NPs from endogenous metabolites of reporter cells, and “D_2_O Sorter” is used to quickly reveal masses of interest. We estimate that any NP with its molecular target present in only a few copies per reporter cell can be detected, which is mainly limited by the sensitivity of MS. This list of metabolites is further curated using Global Natural Product Social (GNPS) molecular networking^28^. This service serves to identify compounds with similar MS2 structures from the pool identified, and screens all the compounds against a database of existing metabolites. This dereplication mitigates the potential for compound/scaffold rediscovery. Following this filtering, molecules are ranked by abundance to prioritize metabolites that may be readily purified and structurally elucidated. Considering minimal metabolic shift between the 0% and 25% D2O culture, a large-scale culture in normal water is expected to produce compounds of interest detected by DABS for purification and structure elucidation using Nuclear Magnetic Resonance (NMR) spectroscopy. It is notable that by chemical extraction of reporting cells, we here primarily focus on non-covalently bound NPs as they are more challenging to be identified through traditional assays and the majority of bioactive NPs likely fall in this category.

Known natural products from microbial extracts were used to test and optimize the DABS workflow. Particularly, all four *Streptomyces* extracts mentioned above were subjected to the DABS workflow using the cervical cancer HeLa cells for reporting. These extracts contain known metabolites (**Figure 2a-d**) that have varying activities and permeabilities in human cancer cells, allowing for a rapid parameterization and validation of the platform workflow. For example, antimycin belongs to the known anti-cancer family of depsipeptides and its binding to and enrichment in HeLa cells have been previously reported using imaging^7^, representing a positive control which should be revealed in our DABS workflow. In contrast, tetracycline, a broad-spectrum antibiotic, lacks the means to appreciably concentrate in mammalian cell lines without the appropriate uptake mechanism^29^; physostigmine, a parasympathomimetic drug, binds covalently to cholinesterase upon transesterification^30^; and asukamycin, an anti-cancer polyketide, acts as a molecular glue via multicovalent attachment^31^. None of them was thus expected to be revealed by DABS. We successfully detected antimycin family of depsipeptides as major binders from *S. albus* extracts via DABS, but did not detect tetracycline, physostigmine, nor asukamycin, an observation consistent with their predicted mode of actions. These results suggested the utility of DABS to reveal non-covalent binders from a mixture and set the stage for discovering new bioactive natural products via DABS.

### Discovery of zillamycin through DABS

Using the four *Streptomyces* extracts, additional deuterated metabolites were also revealed via DABS in addition to antimycins (**Table S1-S4**). Upon dereplication and also considering the abundance of metabolites for subsequent structural elucidation using NMR, we focused on a new metabolite (m/z=468.43, named zillamycin) produced by *S. griseofuscus* (**Figure 3**). *S. griseofuscus* is a well-studied organism that has been explored for usage as a heterologous host as well as in the production of food preservatives^32,33^. In addition to physostigmine, other NPs such as azinomycin a/b and bundlin a/b have been associated with *S. griseofuscus*^34,35^. Zillamycin showed no similarity with known metabolites from *S. griseofuscus* or other strains based on the molecular networking analysis.

For purification and structural elucidation of zillamycin, a four-liter MS-agar culture of *S. griseofuscus* was extracted and purified by size-exclusion chromatography with a Sephadex LH- 20 column packing, followed by reverse-phase chromatography with an SP-C18 packed column and High Performance Liquid Chromatography (HPLC) purification with a Sephadex 250 × 10mm C18 column. zillamycin was isolated as a white amorphous solid with a yield of ∼1 mg, and predicted to have the molecular formula of C_31_H_54_N_3_ based on a proton adduct ion at *m/z* 468.4316 [M+H]^+^ (**Figure 3d** and **Figure S2**). A detailed analysis of the 1D and 2D NMR spectra (^1^H, ^13^C, HSQC, HMBC, and COSY) revealed zillamycin to be a triterpene guanidine with a linear isoprenyl tail of six isoprene units connected to a guanidine moiety via a C-N bond (**Figure 3c, Table S5**, and **Figure S3-S7**).

**Figure 3:**
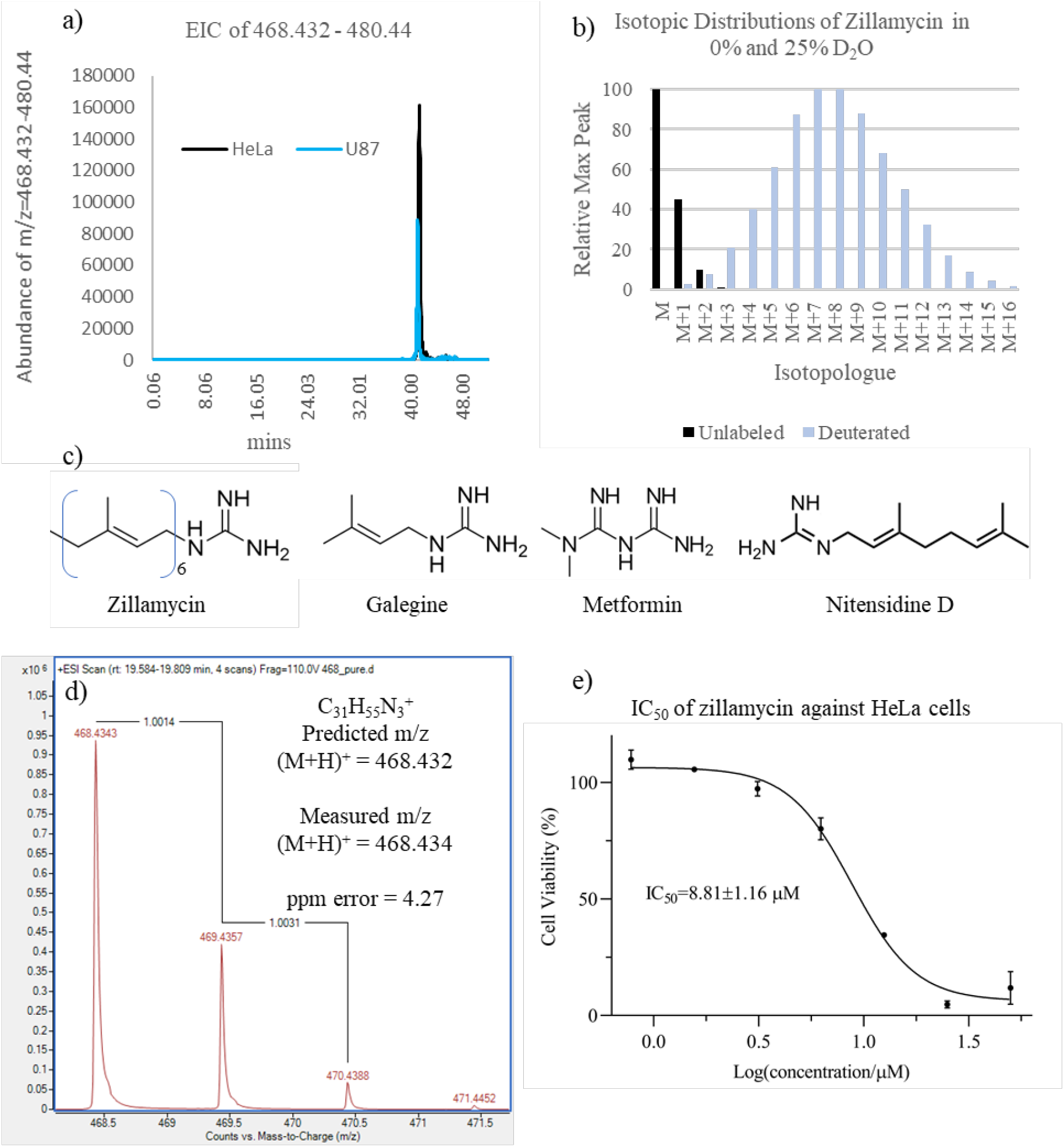
Discovery, structural elucidation, and bioactivities of zillamycin. A metabolite from the extract of *Streptomyces griseofiisctis* with an m/z=468.432 was detected as both deuterated and cell binding, satisfying the DABS criteria, a) Extracted Ion Chromatogram (EIC) of m/z=468.432 and it numerous isotopologues following DABS of *S. griseofitscus* extracts against HeLa cells (black) and U87 cells (blue), b) the unlabeled and deuterated isotopologue distribution of zillamycin. c) Structure of zillamycin and known bioactive compounds with similar structures» d) the Electrospray Ionization (ESI) scan of unlabeled zillamycin and its predicted formula» e) and the IC_50_ of zillamycin against HeLa cells.

The structure of zillamycin confirmed it to be a new natural product never isolated before, however, similar isoprenyl guanidines have been reported as toxic alkaloids from plants. Examples include nitensidine D from the leaves of the South American legume *Pterogyne nitens* and galegine originally isolated from Goat’s rue (*Galega officinalis*) which has led to the development of metformin for diabetes treatment (Figure 3c)^36-38^. Anti-cancer activities have been reported for the known isoprenyl guanidines^39^, and we thus evaluated the cytotoxic activity of Zillamycin against the HeLa cells. Zillamycin showed an anti-cancer activity with an IC_50_ value of 8.81 µM (**Figure 3e**), consistent with the known cytotoxic activity of isoprenyl guanidines.

In addition to the Hela cells, we also assessed the binding capability of zillamycin to other cells, including the brain cancer U87 cells and microbial cells such as the yeast *Candida albicans* and the bacteria *Pseudomonas aeruginosa* and *Streptococcus mutans*. Zillamycin was enriched and revealed from all of these reporting cells (**Figure 3a** and **Figure S8**), suggesting a general binding mechanism. A subsequent cellular localization study with *S. mutans* demonstrated that zillamycin mainly accumulated in the cellular membrane rather than cytosol or cell wall (**Figure S9**), most likely due to its extensive isoprenyl chain. Interestingly, zillamycin demonstrated antimicrobial activities towards selected Gram-positive bacteria with the minimum inhibitory concentration (MIC) values of ∼12.5 µM, but was inactive against the yeast *C. albicans* and the Gram-negative bacterium *P. aeruginosa* up to 50 µM (**Table S6**). The observed different antimicrobial activities of zillamycin suggested that cell binding activities might not necessarily be associated with cytotoxic activities of compounds. This observation was consistent with our previous activity-distribution relationship study of anti-cancer antimycin-type depsipeptides, which demonstrated that the intracellular enrichment and distribution of compounds were driven by their potency and specific protein targets, as well as the lipophilic properties of compounds^16^. In addition to partitioning into membrane fractions, zillamycin may have specific molecular targets in organisms towards which cytotoxicity was observed. We further conclude that to increase the likelihood of bioactive compound discovery, polarity of compounds should be factored in for candidate selection from whole cell binding assays, as polar compounds are less likely to be revealed due to non-specific cell binding and enrichment.

### Conclusion

By pairing the D_2_O metabolome labeling strategy with an untargeted cell binding assay and an LC-MS readout, the DABS workflow provides a unique opportunity to observe interesting bioactive metabolites missed in traditional screenings. This methodology takes advantage of the holistic tracking provided by deuterium, and the broad spectrum of bioactivities identified by cell binding, to produce an assay that provides the means to identify novel natural products with a wide range of bioactive potential. DABS is thus expected to quickly narrow down a list of potentially bioactive natural products from complex mixtures and accelerate new natural product-based drug discovery.

## Methodology

### Deuterium Tagging and Identification

A database including all potential combinations of formulas between 100-1500 Da was generated using the atoms carbon, nitrogen, oxygen, and hydrogen. These atoms are the most common occurring in biomolecules and allow a representative sampling of most natural products. A MATLAB script generated a list of all monoisotopic formulas containing C, H, N, and O between 100-1100 Dalton. This list contained formulas that are not biologically or physically relevant (e.g., CH_90_), due to electron configurations or the sp orbitals available to any given atom. In order to address this, the formula list was curated by applying several of the Seven Golden Rules (SGR), a set of predicted formula distributions based off chemical frequency in a given mass range^40^. The SGR method addresses unlikely molecular formulas using both physical constraints (e.g., balanced charge states) and also uses a database of existing natural products to provide typical configurations of small molecules versus extremely rare formulas. These rules were used to eliminate candidate formulas that were more than three standard deviations away from typical formulas, allowing for a much more representative list of potential molecular formulas possible.

From the curated formula list, the expected isotopic distribution of a given formula was predicted using natural isotopologue frequencies of each atom^41^ with a MATLAB script. These frequencies are constant across all natural products and all naturally occurring molecules. A second script was utilized that predicted the isotopic distribution of a compound that is labeled with an extra neutron from deuterium approximately 6% of the time (**Figures S10-S11**). This number was selected as beneath the lower limit of deuterium incorporation observed in known biomolecules at the concentrations used experimentally (25% D_2_O) (**Figure 2**)

One method that sufficiently differentiated the deuterated and naturally occurring isotopologues was the analysis of the (M+1) and (M+2) peaks relative to the (M) peak. These refer to the base peak with no extra neutrons (M), one extra neutron (M+1), and two extra neutrons (M+2) respectively. The (M) ion as discussed here could represent any number of adducts that occur in ESI LCMS including such as a proton or sodium adduct. Using the isotopic distributions, the data was selected for the local maxima of (M+1) and (M+2) peaks in any given mass range. For small molecules, the ratios of these higher peaks over the base peak are typically quite small. In contrast, for deuterated small molecules, the ratios are typically quite large (shown in red, **Figure S11**). A curve representing the maximum isotopic distributions observed in naturally occurring samples is shown as the green line. Statistics regarding the frequency of deuterated samples exceeding this curvature were predicted to examine deuterium’s detection capabilities. Combining both data sets, molecules that are deuterated supersede the maxima line of both sets 93.5% of the time. This means that with 93.5% of molecules deuterated to 6%, using this criterion allows a high level assurance that the molecule has been artificially tagged and virtually eliminates false negatives (**Figure S10**). Compounds failing to pass this requirement are typically highly unsaturated molecules with low hydrogen and high oxygen counts. Curiously, a significant deviation in carbon formula abundance was also illustrated, with low carbon counts prohibiting detection. This is expected to be due to the low carbon count constraining the max number of hydrogens based off SGR predictions.

Mass spectrometry data was collected using Mass Hunter Data Acquisition. These files were exported into the preprocessing software MS-Dial. MS-Dial performs retention time corrections and collapses centroided data into single peaks per ion (peak picking). A MATLAB script “D_2_O Sorter” was written to predict whether a given small molecule had been labeled. The MATLAB code for this program is included in the SI. This script’s input included the peak picked data from MS-Dial. It sorts for peaks expected to be isotopes of each other. Significant retention time drift was allowed between samples due to hydrophobic secondary isotope effects induced by deuterium. Each addition of a deuterium atom slightly increased the molecules polarity. These compiled peak lists were then screened with their isotopologue distributions compared to the natural maxima predictions of the given mass region. Depending on whether they met the (M+2) and (M+1) relative height requirements, the peaks were sorted into respective “Labeled” and “Unlabeled” matrixes. These matrixes provide mass, retention time, and abundance data. These results can then be manually observed and referenced back to the original Mass Hunter data to determine which masses correspond to the labeled compounds of interest.

### Cell Binding

Strains known to produce a number of known natural products were selected for validation. This list comprised of a variety of *Streptomyces* with a prominent and well characterized metabolite present. Although *Streptomyces* produce dozens of uncharacterized metabolites, these known small molecules provide a proof of concept for labeling.

The various *Streptomyces* strains were cultured using Mannitol Soya (MS) media on agar for 10 days at 30 C. The strains were extracted 1:1 with organic solvent using 95:5 ethyl acetate:methanol and incubated for 30 minutes during extraction. The extracts were dried and resuspended at 10x in DMSO.

For mammalian cells, cultures are grown to 90% confluency adhered to a 12 well plate in DMEM+ 5% FBS media. The cell culture extracts are added at a 1 mL extract equivalent / 1 mL mammalian cell culture equivalent and allowed to incubate for four hours at 37C. After four hours, the cells are gently washed with PBS three times. After washing, the cells are resuspended in 1 mL of PBS and vigorously pipetted to detach the cells. These detached cells are extracted 1:1 with 95:5 ethyl acetate:methanol. The resulting extract was dried down and resuspended at 10x in methanol. 100uL human cell culture equivalent was injected onto the LCMS.

Microbial cells are grown to pre-stationary phase. The cells are concentrated to 3x to 1 mL of cell culture. The cells are centrifuged during wash steps to allow supernatant separation from the pellet. All other steps are the same as those performed on mammalian cells.

### Purification of zillamycin

4Ls of *S. griseofuscus* was grown on MS-agar plates as lawns for 12 days. Upon the last day, the plates were extracted with 80:20 ethyl acetate:methanol. This organic phase was centrifuged to collapse the interphase, and the organic phase was separated from the remaining components. This extract was dried down using a rotovap and resuspended in 5 mLs methanol. The extract was quite oily and experienced phase separation at lower volumes of methanol.

This extracted sample was run through a packed LH-20 size exclusion column, with fractions being tested by LCMS for presence of zillamycin. Appendix F.1C shows the elution of the compound as detected through fractions tested on a mass spectrometer. The fraction containing zillamycin was dried and resuspended in 1mL of methanol. These contents were separated using an SP-C18 packed column. The column was washed with acetonitrile + 1% TFA and methanol, before eluting the compound with methanol + 1% TFA.

This fraction was concentrated to 1 mL of methanol and injected onto the HPLC in 100uL injections. Specifics of the HPLC method are detailed in appendix F.1B. The fraction containing eluted at 20 mins with a UV maximum of 208nm (though zillamycin has a fairly weak UV single).

### Bioactivity Assays

#### HeLa SRB IC50 Assays

HeLa cells were dosed with zillamycin up to 50 uM at 1% DMSO using the standard SRB protocol^42^ (Figure 3e).

#### Antimicrobial Assays

Various microbial strains were grown in brain heart infusion (BHI) media to an OD=1. The cells were diluted (2000x) into a 200uL 96 well plate to an OD=0.0005. The wells were dosed with 2 uL DMSO (including blanks and positive control antibiotics) in duplicate. The 96 well plate was incubated at 37C in a plate reader for 36 hours. Graphical results of cell growth are shown in Appendix C.

#### Compound Localization

The subcellular fractionation protocol was adapted from Ahn. et al^43^. The resulting fractions include a sample of the unlysed cellular components, supernatant, cell wall, cell membrane, and the cytosol. With dosed zillamycin, significant accumulation was seen in the cellular membrane, though some was also present in the cytosol (**Figure S8**).

## Supporting information

DABS_Supporting_Information

## Acknowledgements

This research was financially supported by grants to W.Z. from Southern Marine Science and Engineering Guangdong Laboratory (Guangzhou) (SMSEGL20SC02).

The authors declare no competing financial interest or conflict of interest.

## References

1) Newman, D. J.; Cragg, G. M. Natural Products as Sources of New Drugs over the Nearly Four Decades from 01/1981 to 09/2019. Journal of Natural Products 2020, 83 (3).

(2) Chen, Y.; Garcia de Lomana, M.; Friedrich, N.-O.; Kirchmair, J. Characterization of the Chemical Space of Known and Readily Obtainable Natural Products. Journal of Chemical Information and Modeling 2018, 58 (8), 1518–1532.

(3) Nelson, A.; Karageorgis, G. Natural Product-Informed Exploration of Chemical Space to Enable Bioactive Molecular Discovery. RSC Medicinal Chemistry 2021, 12 (3), 353–362.

(4) Rosén, J.; Gottfries, J.; Muresan, S.; Backlund, A.; Oprea, T. I. Novel Chemical Space Exploration via Natural Products. Journal of Medicinal Chemistry 2009, 52 (7), 1953–1962.

(5) Conly, J.; Johnston, B. Where Are All the New Antibiotics? The New Antibiotic Paradox. Canadian Journal of Infectious Diseases and Medical Microbiology 2005, 16 (3), 159–160.

(6) Chang, J.; Kwon, H. J. Discovery of Novel Drug Targets and Their Functions Using Phenotypic Screening of Natural Products. Journal of Industrial Microbiology and Biotechnology 2016, 43 (2-3), 221–231.

(7) Scherlach, K.; Hertweck, C. Mining and Unearthing Hidden Biosynthetic Potential. Nature Communications 2021, 12 (1), 3864.

(8) Bauman, K. D.; Butler, K. S.; Moore, B. S.; Chekan, J. R. Genome Mining Methods to Discover Bioactive Natural Products. Natural Product Reports 2021, 38 (11), 2100–2129.

(9) Deutsch, J. M.; Mandelare-Ruiz, P.; Yang, Y.; Foster, G.; Routhu, A.; Houk, J.; De La Flor, Y. T.; Ushijima, B.; Meyer, J. L.; Paul, V. J.; Garg, N. Metabolomics Approaches to Dereplicate Natural Products from Coral-Derived Bioactive Bacteria. Journal of Natural Products 2022.

(10) Demarque, D. P.; Dusi, R. G.; de Sousa, F. D. M.; Grossi, S. M.; Silvério, M. R. S.; Lopes, N. P.; Espindola, L. S. Mass Spectrometry-Based Metabolomics Approach in the Isolation of Bioactive Natural Products. Scientific Reports 2020, 10 (1).

(11) Caesar, L. K.; Montaser, R.; Keller, N. P.; Kelleher, N. L. Metabolomics and Genomics in Natural Products Research: Complementary Tools for Targeting New Chemical Entities. Natural Product Reports 2021, 38 (11), 2041–2065.

(12) Petras, D.; Phelan, V. V.; Acharya, D.; Allen, A. E.; Aron, A. T.; Bandeira, N.; Bowen, B. P.; Belle-Oudry, D.; Boecker, S.; Cummings, D. A.; Deutsch, J. M.; Fahy, E.; Garg, N.; Gregor, R.; Handelsman, J.; Navarro-Hoyos, M.; Jarmusch, A. K.; Jarmusch, S. A.; Louie, K.; Maloney, K. N. GNPS Dashboard: Collaborative Exploration of Mass Spectrometry Data in the Web Browser. Nature Methods 2022, 19 (2), 134–136.

(13) Seyedsayamdost, M. R.; Clardy, J. Natural Products and Synthetic Biology. ACS Synthetic Biology 2014, 3 (10), 745–747.

(14) Jamieson, C. S.; Misa, J.; Tang, Y.; Billingsley, J. M. Biosynthesis and Synthetic Biology of Psychoactive Natural Products. Chemical Society Reviews 2021, 50 (12), 6950–7008.

(15) Ory, L.; Nazih, E.-H.; Daoud, S.; Mocquard, J.; Bourjot, M.; Margueritte, L.; Delsuc, M.-A.; Bard, J.-M.; Pouchus, Y. F.; Bertrand, S.; Roullier, C. Targeting Bioactive Compounds in Natural Extracts - Development of a Comprehensive Workflow Combining Chemical and Biological Data. Analytica Chemical Acta 2019, 1070, 29–42.

(16) Seidel, J., Miao, Y., Porterfield, W., Cai, W., Zhu, X., Kim, S., Hu, F., Bhattarai-Kline, S., Min, W., Zhang, W. “Structure-activity-distribution relationship study of anti-cancer antimycin-type depsipetides.” Chem. Commun. 2019, 55, 9379–9382.

(17) Mukne, A.; Momin, M.; Betkar, P.; Rane, T.; Valecha, S. Cell-Based Assays in Natural Product-Based Drug Discovery. Evidence Based Validation of Traditional Medicines 2021, 211– 248.

(18) Gabriel, J.; Höfner, G.; Wanner, K. T. A Library Screening Strategy Combining the Concepts of MS Binding Assays and Affinity Selection Mass Spectrometry. Frontiers in Chemistry 2019, 7.

(19) Rinkel, J.; Dickschat, J. S. Recent Highlights in Biosynthesis Research Using Stable Isotopes. Beilstein J. Org. Chem. 2015, 11, 2493–2508.

(20) Weindl, D.; Cordes, T.; Battello, N.; Sapcariu, S. C.; Dong, X.; Wegner, A.; Hiller, K. Bridging the Gap between Non-Targeted Stable Isotope Labeling and Metabolic Flux Analysis. Cancer Metab. 2016 41 2016, 4 (1), 1–14.

(21) Palaniappan, K. K.; Pitcher, A. A.; Smart, B. P.; Spiciarich, D. R.; Iavarone, A. T.; Bertozzi, C. R. Isotopic Signature Transfer and Mass Pattern Prediction (IsoStamp): An Enabling Technique for Chemically-Directed Proteomics. ACS Chem. Biol. 2011, 6 (8), 829–836.

(22) Llufrio, E. M.; Cho, K.; Patti, G. J. Systems-Level Analysis of Isotopic Labeling in Untargeted Metabolomic Data by X13CMS. Nature Protocols 2019, 14 (7), 1970–1990.

(23) Kushner, D. J.; Baker, A.; Dunstall, T. G. Pharmacological Uses and Perspectives of Heavy Water and Deuterated Compounds. Can. J. Physiol. Pharmacol. 1999, 77 (2), 79–88.

(24) Berry, D.; Mader, E.; Lee, T. K.; Woebken, D.; Wang, Y.; Zhu, D.; Palatinszky, M.; Schintlmeister, A.; Schmid, M. C.; Hanson, B. T.; et al. Tracking Heavy Water (D2O) Incorporation for Identifying and Sorting Active Microbial Cells. Proc. Natl. Acad. Sci. U. S. A. 2015, 112 (2), E194–E203.

(25) Xu, F.; Nazari, B.; Moon, K.; Bushin, L. B.; Seyedsayamdost, M. R. Discovery of a Cryptic Antifungal Compound from Streptomyces Albus J1074 Using High-Throughput Elicitor Screens. Journal of the American Chemical Society 2017, 139 (27), 9203–9212.

(26) Nguyen, T. B.; Kitani, S.; Shimma, S.; Nihira, T. Butenolides From Streptomyces Albus J1074 Act as External Signals to Stimulate Avermectin Production In Streptomyces Avermitilis. Applied and Environmental Microbiology 2018, 84 (9).

(27) Turowski, M.; Yamakawa, N.; Meller, J.; Kimata, K.; Ikegami, T.; Hosoya, K.; Tanaka, N.; Thornton, E. R. Deuterium Isotope Effects on Hydrophobic Interactions: The Importance of Dispersion Interactions in the Hydrophobic Phase. Journal of the American Chemical Society 2003, 125 (45), 13836–13849.

(28) Wang, M.; Carver, J. J.; Phelan, V. V.; Sanchez, L. M.; Garg, N.; Peng, Y.; Nguyen, D. D.; Watrous, J.; Kapono, C. A.; Luzzatto-Knaan, T.; et al. Sharing and Community Curation of Mass Spectrometry Data with Global Natural Products Social Molecular Networking. Nat. Biotechnol. 2016, 34 (8), 828–837.

(29) Chopra, I.; Roberts, M. Tetracycline Antibiotics: Mode of Action, Applications, Molecular Biology, and Epidemiology of Bacterial Resistance. Microbiology and Molecular Biology Reviews 2001, 65 (2), 232–260.

(30) Triggle, D. J.; Mitchell, J. M.; Filler, R. The Pharmacology of Physostigmine. CNS Drug Reviews 1998, 4 (2), 87–136.

(31) Geiger, T. M.; Schäfer, S. C.; Dreizler, J. K.; Walz, M.; Hausch, F. Clues to Molecular Glues. Current Research in Chemical Biology 2022, 2, 100018.

(32) Gren, T.; Whitford, C. M.; Mohite, O. S.; Jørgensen, T. S.; Kontou, E. E.; Nielsen, J. B.; Lee, S. Y.; Weber, T. Characterization and Engineering of Streptomyces Griseofuscus DSM 40191 as a Potential Host for Heterologous Expression of Biosynthetic Gene Clusters. Scientific Reports 2021, 11 (1), 18301.

(33) Li, S.; Tang, L.; Chen, X.; Liao, L.; Li, F.; Mao, Z. Isolation and Characterization of a Novel ε-Poly-l-Lysine Producing Strain: Streptomyces Griseofuscus. Journal of Industrial Microbiology & Biotechnology 2010, 38 (4), 557–563.

(34) Ishizeki, S.; Ohtsuka, M.; Irinoda, K.; Kukita, K.-I.; Nagaoka, K.; Nakashima, T. Azinomycins a and B, New Antitumor Antibiotics. III. Antitumor Activity. The Journal of Antibiotics 1987, 40 (1), 60–65.

(35) Sakamoto, J. M. J.; Kondō, S.-I.; Yumoto, H.; Arishima, M. Bundlins a and B, Two Antibiotics Produced by Streptomyces Griseofuscus Nov. Sp. The Journal of Antibiotics, Series A 1962, 15 (2), 98–102.

(36) Bailey, C. J.; Turner, R. C. Metformin. New England Journal of Medicine 1996, 334 (9), 574– 579.

(37) Mooney, M. H.; Fogarty, S.; Stevenson, C.; Gallagher, A. M.; Palit, P.; Hawley, S. A.; Hardie, D. G.; Coxon, G. D.; Waigh, R. D.; Tate, R. J.; et al. Mechanisms Underlying the Metabolic Actions of Galegine That Contribute to Weight Loss in Mice. Br. J. Pharmacol. 2008, 153, 1669– 1677.

(38) Regasini, L. O.; Castro-Gamboa, I.; Silva, D. H. S.; Furlan, M.; Barreiro, E. J.; Ferreira, P. M. P.; Pessoa, C.; Lotufo, L. V. C.; De Moraes, M. O.; Young, M. C. M.; et al. Cytotoxic Guanidine Alkaloids from Pterogyne Nitens. J. Nat. Prod. 2009.

(39) Wang, H. X.; Ng, T. B. Natural Products with Hypoglycemic, Hypotensive, Hypocholesterolemic, Antiatherosclerotic and Antithrombotic Activities. Life Sci. 1999, 65 (25), 2663–2677.

(40) Kind, T.; Fiehn, O. Seven Golden Rules for Heuristic Filtering of Molecular Formulas Obtained by Accurate Mass Spectrometry. BMC Bioinformatics 2007, 8, 1–20.

(41) Mass Spectrometry: Isotope Effects https://chem.libretexts.org/Bookshelves/Analytical_Chemistry/Supplemental_Modules_(Analytical_Chemistry)/Instrumental_Analysis/Mass_Spectrometry/Mass_Spectrometry%3A_Isotope_Effects (accessed 2023 -01 -27).).

(42) SRB Assay / Sulforhodamine B Assay Kit (ab235935) | Abcam https://www.abcam.com/srb-assay--sulforhodamine-b-assay-kit-ab235935.html (accessed Apr 27, 2022).

(43) Ahn, S. J.; Kaspar, J.; Kim, J. N.; Seaton, K.; Burne, R. A. Discovery of Novel Peptides Regulating Competence Development in Streptococcus Mutans. J. Bacteriol. 2014, 196 (21), 3735.

