## Supplementary material for "Bioactive Natural Product Discovery via Deuterium Adduct Bioactivity Screening": DABS_Supporting_Information

This research was financially supported by grants to W.Z. from Southern Marine Science and Engineering Guangdong Laboratory (Guangzhou) (SMSEGL20SC02).

### Figures:

FIGURE S8: OVERLAYED MASS SPECTRA OF ZILLAMYCIN FOLLOWING DABS THROUGH VARIOUS MICROBIAL SPECIES INCLUDING CANDIDA ALBICANS (RED), PSEUDOMONAS AERUGINOSA (GREY), STREPTOCOCCUS MUTANS (YELLOW), AND A BLANK (BLUE). THE Y-AXIS IS SCALE TO THE MAX OF THE BLANK TO PROVIDE A SIGNAL TO NOISE RATIO. ALL THREE STRAINS SHOWED SIGNIFICANT BINDING TO THE SMALL MOLECULE. EACH CULTURE WAS GROWN OVERNIGHT IN BHI. THE FOLLOWING DAY, 1 ML SAMPLES WERE ALIQUOTED FROM THE OVERNIGHT CULTURES AND WERE INOCULATED WITH 5 MG/L ZILLAMYCIN. THESE SAMPLES WERE CULTURED FOR 4 HOURS AT 37C. FOLLOWING CULTURING, EACH STRAIN WAS SPUN DOWN AND WASHED WITH FRESH PBS FOR 3X TIMES. FOLLOWING THE THIRD WASH, THESE SAMPLES WERE EXTRACTED IN METHANOL AND INJECTED ONTO THE MASS SPECTROMETER. ........13
FIGURE S9: SUBCELLULAR MEMBRANE FRACTIONATION OF STREPTOCOCCUS MUTANS INCUBATED FOR 4 HOURS WITH ZILLAMYCIN. RESULTS INDICATE THAT THE COMPOUND SIGNIFICANTLY LOCALIZES WITH THE CELLULAR MEMBRANE........14
FIGURE S10: PROCESS FOR GENERATING A COMPREHENSIVE MASS AND FORMULA LIST WITH ISOTOPOLOGUE ABUNDANCES. A COMBINATORIAL LIST OF THE POSSIBLE FORMULAS CONTAINING CARBON, HYDROGEN, NITROGEN, AND OXYGEN FROM MASSES 100-1100 DALTON WAS GENERATED. THIS LIST WAS PAIRED DOWN FOR “REALISTIC” FORMULAS BY REFERENCING RULES SET OUT IN THE “SEVEN GOLDEN RULES” PAPER. SPECIFICALLY, THE NUMBER OF EACH ELEMENT ALLOWED IN A MASS RANGE, THE FOLLOWING OF LEWIS RULES, H/C RATIOS, H/N, AND H/O RATIOS WERE ALL MODULATED TO REFLECT WITHIN THREE STANDARD DEVIATIONS OF EXISTING NATURAL PRODUCTS. FROM THE REMAINING FORMULAS, BOTH NATURAL AND DEUTERATED (6%) ISOTOPIC ABUNDANCES WERE CALCULATED........15
FIGURE S11: RELATIVE MASS DIFFERENCE BETWEEN A) (M+1)/M AND MASS OR B) (M+2)/M AND MASS. PLOTTED IN BOTH GRAPHS ARE BOTH THE NATURAL DISTRIBUTION (BLUE STAR) AND THE 6% DEUTERATED DISTRIBUTION (RED STAR) OF THE ISOTOPOLOGUES. THE PLOTTED GREEN LINE SHOWS THE MAXIMA OF THE NATURAL

Table S1: Most abundant mass peaks selected for *Streptomyces albus* metabolites bound to HeLa cells.

| Top Masses Detected (m/z) | Retention Time (mins) | Cumulative Isotopologue Abundance | Relative M+1 / M | Relative M+2 / M |
|---|---|---|---|---|
| 537.2 | 40.6 | 4.071E+05 | 0.47 | 0.11 |
| 549.2 | 42.3 | 3.896E+05 | 0.48 | 0.11 |
| 265.1 | 40.6 | 2.078E+05 | 0.26 | 0.03 |
| 521.2 | 41.2 | 1.723E+05 | 0.46 | 0.10 |
| 1568.8 | 44.5 | 1.491E+05 | 1.28 | 0.77 |
| 537.2 | 40.1 | 1.359E+05 | 0.47 | 0.11 |
| 265.1 | 42.2 | 1.264E+05 | 0.26 | 0.03 |
| 564.2 | 42.5 | 1.206E+05 | 0.49 | 0.12 |
| 784.7 | 27.3 | 1.064E+05 | 0.66 | 0.21 |
| 121.1 | 2.1 | 1.010E+05 | 0.14 | 0.01 |

Table S2: Most abundant mass peaks selected for *Streptomyces aureofaciens* metabolites bound to HeLa cells.

| Top Masses Detected (m/z) | Retention Time (mins) | Cumulative Isotopologue Abundance | Relative M+1 / M | Relative M+2 / M |
|---|---|---|---|---|
| 536.2 | 50.2 | 1.825E+05 | 0.47 | 0.11 |
| 760.2 | 44.1 | 1.578E+05 | 0.64 | 0.20 |
| 550.5 | 40.2 | 5.210E+04 | 0.48 | 0.11 |
| 371.1 | 43.7 | 4.533E+04 | 0.34 | 0.06 |
| 403.4 | 35.0 | 4.324E+04 | 0.36 | 0.06 |
| 945.2 | 49.7 | 4.291E+04 | 0.79 | 0.30 |
| 1295.4 | 48.1 | 4.118E+04 | 1.06 | 0.54 |
| 473.5 | 39.2 | 3.527E+04 | 0.42 | 0.08 |
| 826.3 | 28.6 | 2.430E+04 | 0.69 | 0.23 |

Table S3: Most abundant mass peaks selected for *Streptomyces griseofuscus* metabolites bound to HeLa cells.

| Top Masses Detected (m/z) | Retention Time (mins) | Cumulative Isotopologue Abundance | Relative M+1 / M | Relative M+2 / M |
|---|---|---|---|---|
| 468.4* | 41.4 | 6.036E+06 | 2.53 | 2.57 |
| 478.5 | 41.2 | 2.624E+06 | 1.49 | 1.5 |
| 342.2 | 19.5 | 2.178E+06 | 4.42 | 3.62 |
| 380.2 | 19.9 | 1.707E+06 | 0.64 | 15.3 |
| 882.5 | 22.8 | 1.106E+06 | 1.96 | 0.41 |
| 929.5 | 21.4 | 1.092E+06 | 2.17 | 0.32 |

*zillamycin

Table S4: Most abundant mass peaks selected for *Streptomyces nodosus* metabolites bound to HeLa cells.

| Top Masses Detected (m/z) | Retention Time (mins) | Cumulative Isotopologue Abundance | Relative M+1 / M | Relative M+2 / M |
|---|---|---|---|---|
| 442.1 | 24.8 | 9.866E+04 | 0.39 | 0.08 |
| 1020.2 | 48.5 | 8.206E+04 | 0.85 | 0.34 |
| 442.1 | 26.1 | 7.948E+04 | 0.39 | 0.08 |
| 998.3 | 48.2 | 6.405E+04 | 0.83 | 0.33 |
| 142.0 | 2.6 | 5.465E+04 | 0.16 | 0.01 |
| 758.2 | 44.3 | 5.026E+04 | 0.64 | 0.20 |
| 532.3 | 39.5 | 3.702E+04 | 0.46 | 0.10 |
| 1389.3 | 48.6 | 2.859E+04 | 1.13 | 0.61 |
| 1315.3 | 44.9 | 2.821E+04 | 1.08 | 0.55 |
| 706.5 | 47.9 | 2.661E+04 | 0.60 | 0.17 |

Table S5: The NMR Data of zillamycin in DMSO-$d_6$

| Position | $\delta_H$ (J in Hz) | $\delta_C$ (C type) | Position | $\delta_H$ (J in Hz) | $\delta_C$ (C type) |
|---|---|---|---|---|---|
| 1 | 3.71 dd 6.7, 5.2 | 39.8, $CH_2$ | 17 | 2.02 m | 26.0, $CH_2$ |
| 2 | 5.18 t 6.7 | 118.6, CH | 18 | 5.09 m | 124.0, CH |
| 3 | | 139.7, C | 19 | | 134.3, C |
| 4 | 1.99 m | 38.8, $CH_2$ | 20 | 1.93 m | 38.9, $CH_2$ |
| 5 | 2.05 m | 25.9, $CH_2$ | 21 | 2.02 m | 26.2, $CH_2$ |
| 6 | 5.09 m | 123.6, CH | 22 | 5.07 m | 124.1, CH |
| 7 | | 134.7, C | 23 | | 130.6, C |
| 8 | 1.93 m | 39.2, $CH_2$ | 24 | 1.63 s | 25.5, $CH_3$ |
| 9 | 2.02 m | 26.0, $CH_2$ | 25 | 1.64 s | 16.1, $CH_3$ |
| 10 | 5.07 m | 123.9, CH | 26 | 1.56 s | 15.8, $CH_3$ |
| 11 | | 134.2, C | 27 | 1.55 s | 15.8, $CH_3$ |
| 12 | 1.93 m | 39.1, $CH_2$ | 28 | 1.55 s | 15.8, $CH_3$ |
| 13 | 2.02 m | 26.0, $CH_2$ | 29 | 1.55 s | 15.8, $CH_3$ |
| 14 | 5.09 m | 123.9, CH | 30 | 1.55 s | 17.5, $CH_3$ |
| 15 | | 134.4, C | 31 | | 156.6, C |
| 16 | 1.93 m | 39.2, $CH_2$ | -NH- | 7.50, t 5.2 | |

Table S6: MICs of zillamycin against a variety of microbial cells.

| Species and Strain | Zillamycin MIC (uM) |
|---|---|
| *Pseudomonas aeruginosa* PAO1 | >50 uM |
| *Candida albicans* ATCC 10231 | >50 uM |
| *Bacillus subtilus* 168 | 12.5 uM |
| *Streptococcus salivarius* K12 | 12.5 uM |
| *Streptococcus mutans* S1B | 12.5 uM |
| *Streptomyces albus* J1074 | 12.5 uM |

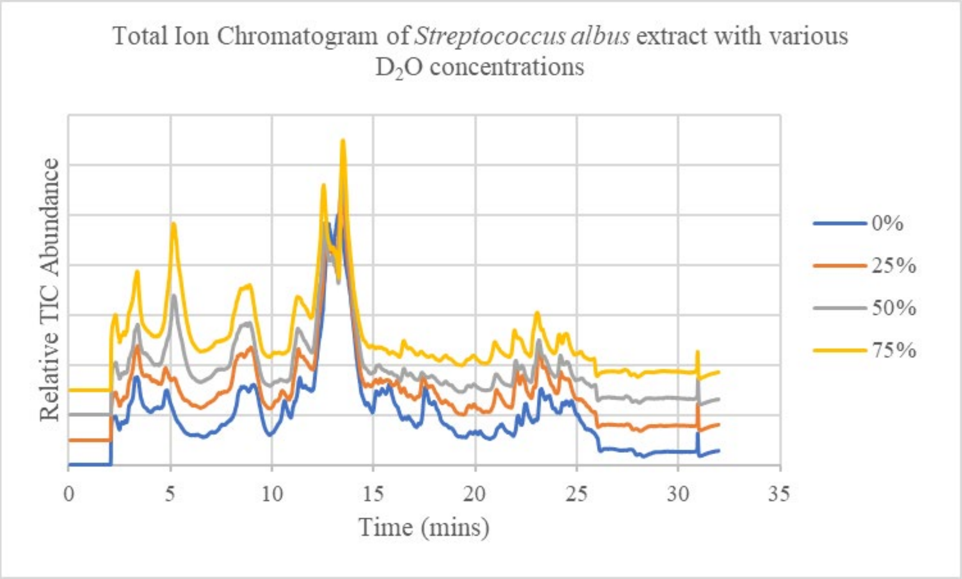

Figure S1: Total Ion Chromatogram of *Streptococcus albus* extract with various $D_2O$ concentrations. Plots are displaced from each other by 10% in increasing $D_2O$ concentration for visibility

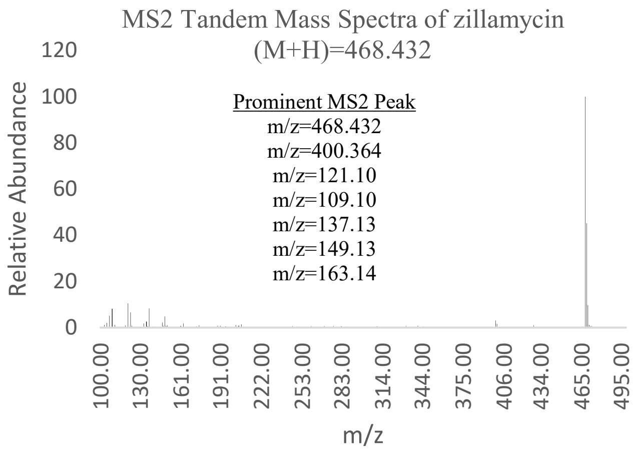

Figure S2: the MS2 spectra of zillamycin

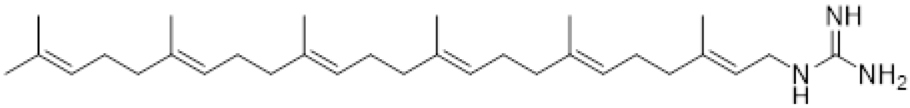

zillamycin ([1]H NMR, DMSO-$d_6$ at 900 MHz)

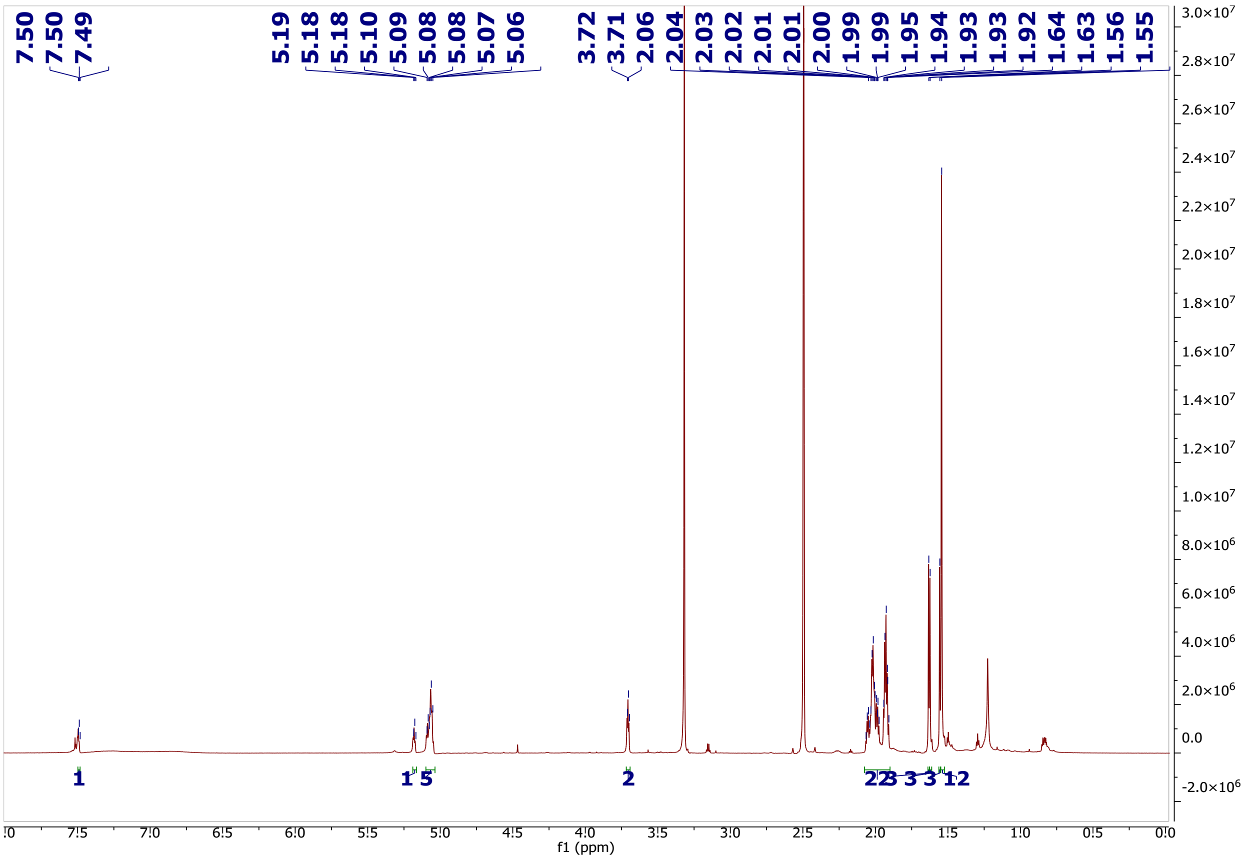

Figure S3: $^{1}$H NMR spectrum of zillamycin

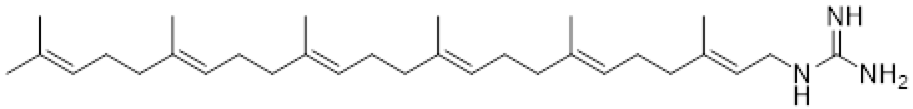

zillamycin ([1]H NMR, DMSO-$d_6$ at 900 MHz)

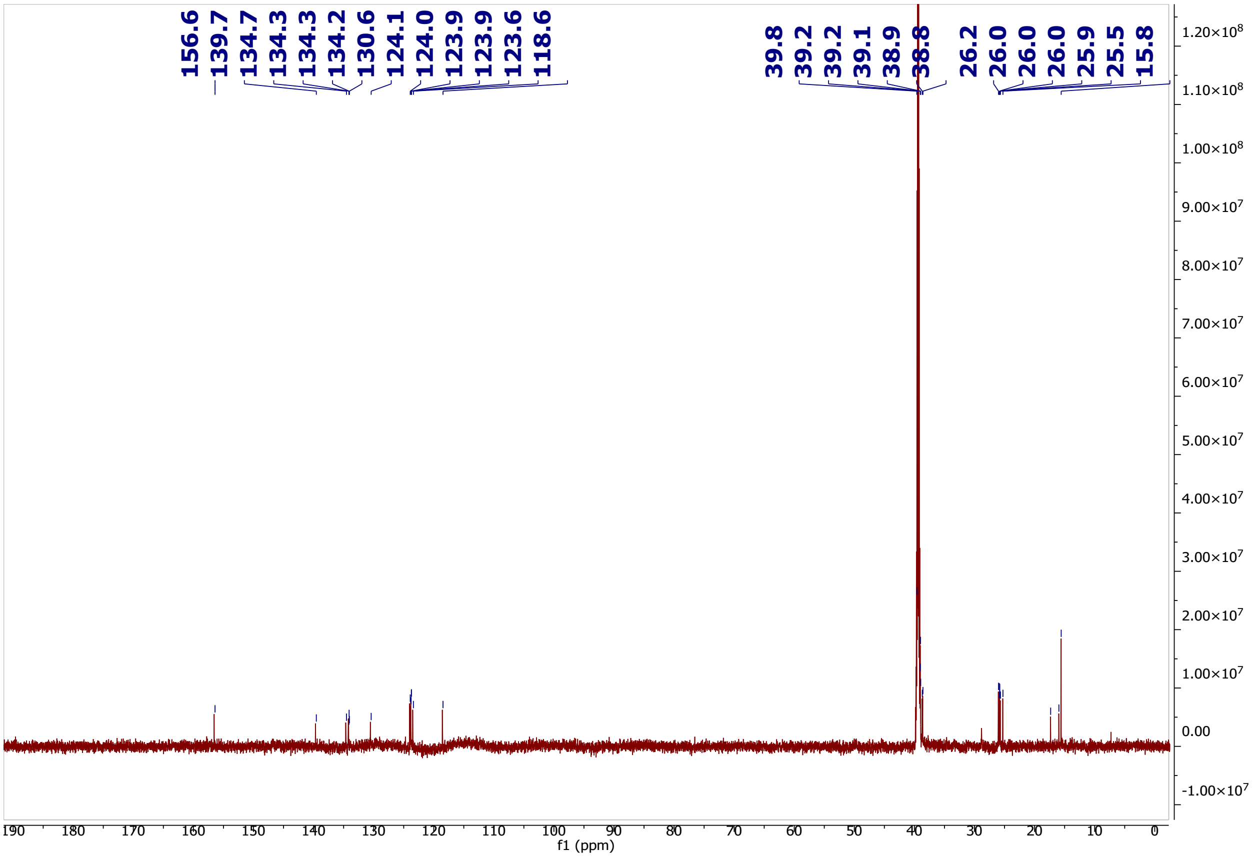

Figure S4: $^{13}C$ NMR spectrum of zillamycin

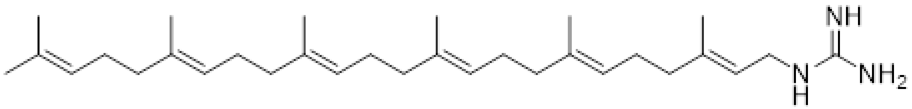

zillamycin ($^1H$-$^1H$ COSY, DMSO-$d_6$ at 900 MHz)

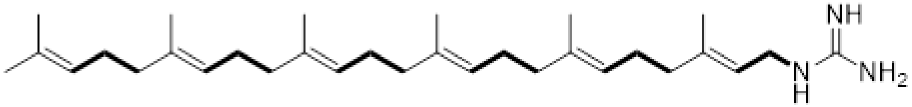

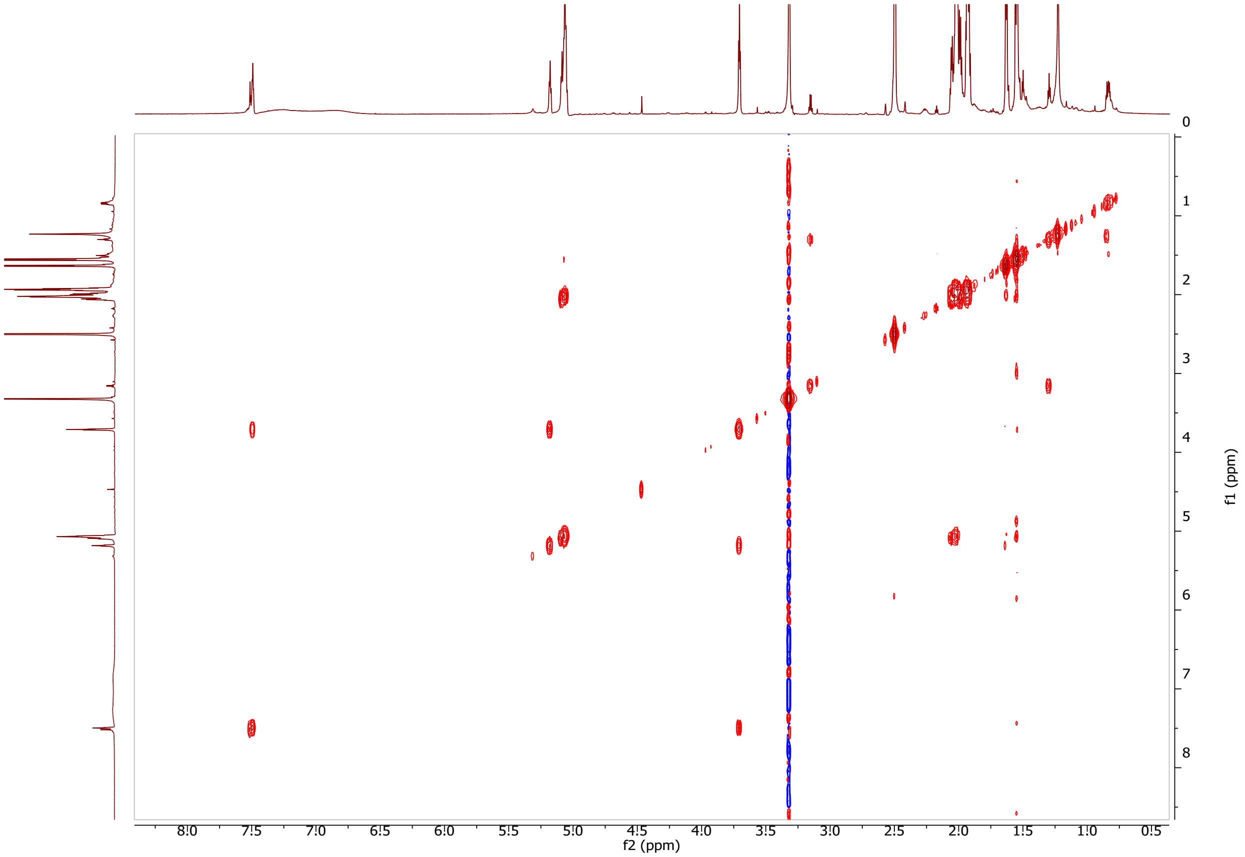

Figure S5: $^1H$-$^1H$ COSY spectrum of zillamycin

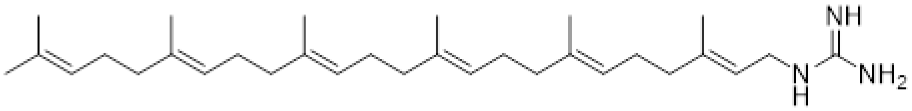

zillamycin ($^1$H-$^{13}$C HSQC, DMSO-$d_6$ at 900 MHz)

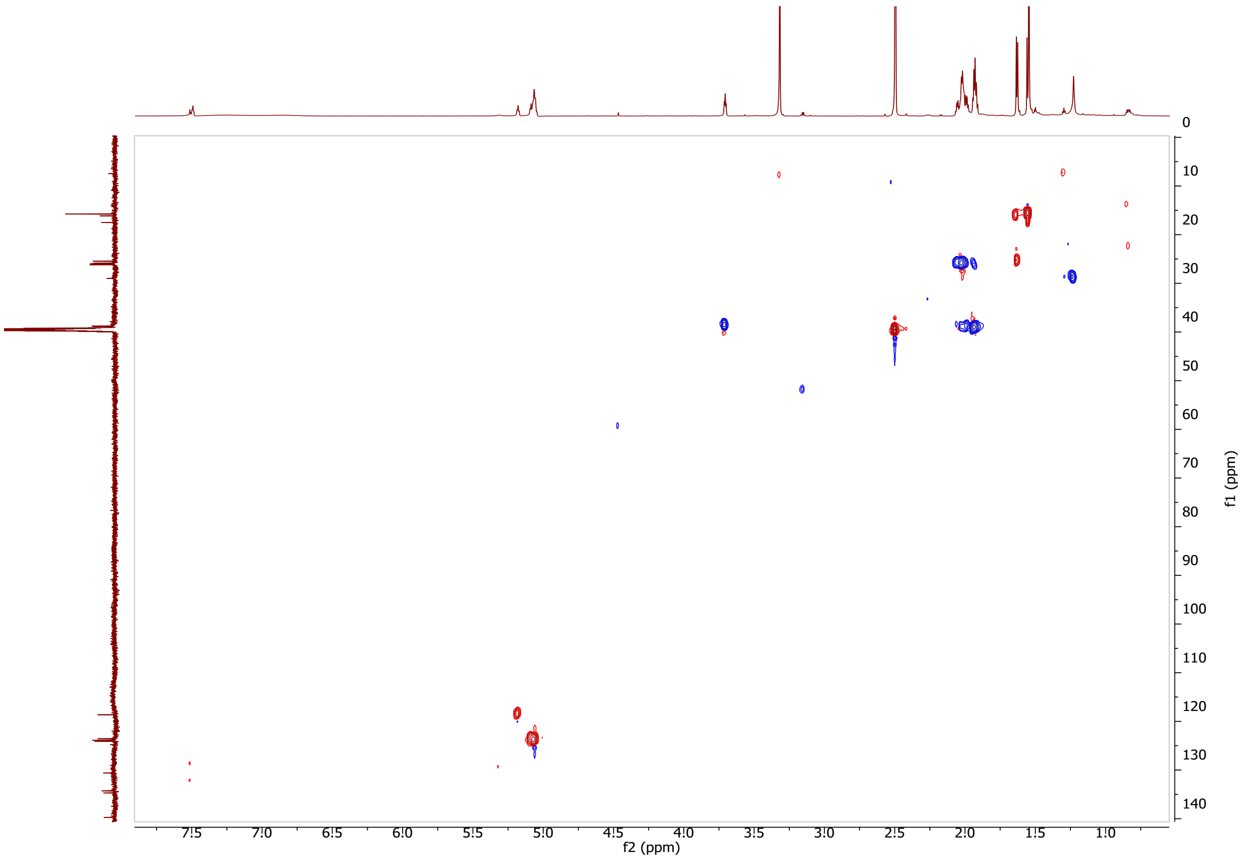

Figure S6: $^1$H-$^{13}$C HSQC spectrum of zillamycin.

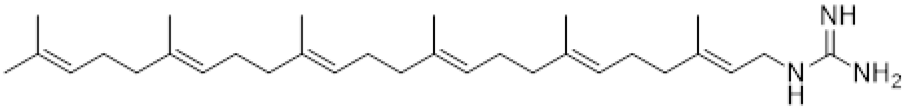

zillamycin ($^{1}H$-$^{1}H$ COSY, DMSO-$d_6$ at 900 MHz)

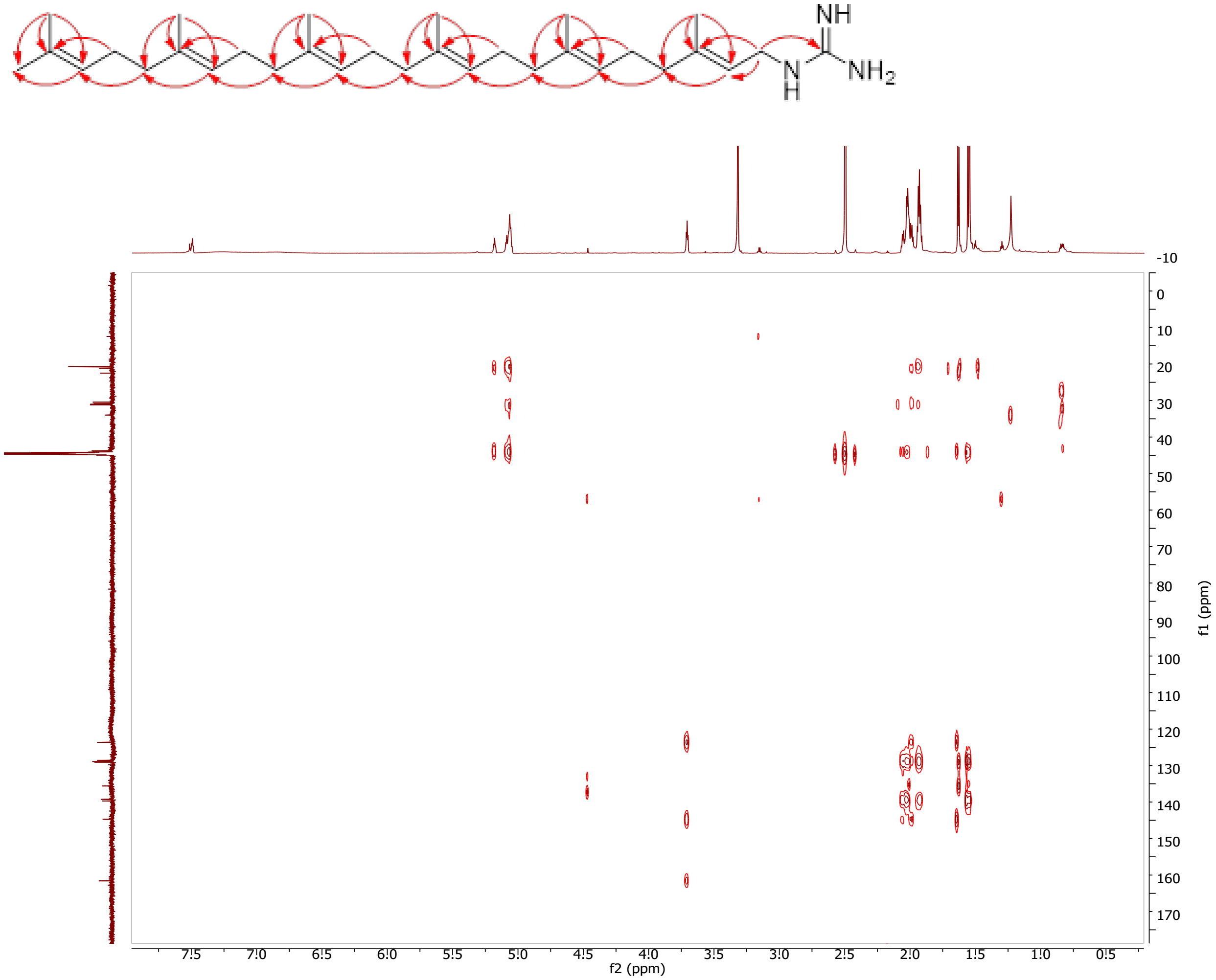

Figure S7: $^{1}H$-$^{13}C$ HMBC spectrum of zillamycin

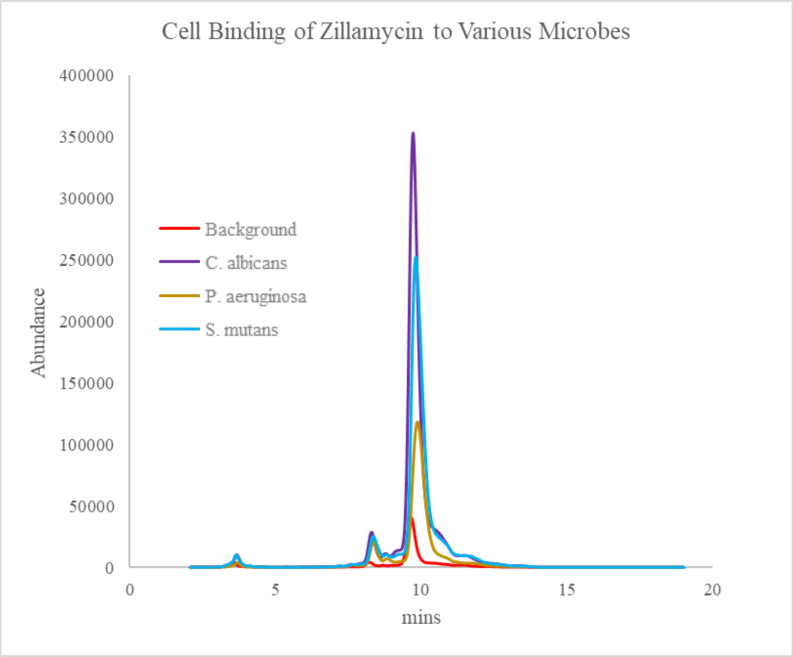

Figure S8: Overlayed mass spectra of zillamycin following DABS through various microbial species including *Candida albicans* (purple), *Pseudomonas aeruginosa* (yellow), *Streptococcus mutans* (blue), and a background (orange). Residual zillamycin is observed following the series wash steps in the background sample, likely due to compound binding to the microcentrifuge tubes. The y-axis is scale to the max of the blank to provide a signal to noise ratio. All three strains showed significant binding to the small molecule. Each culture was grown overnight in BHI reaching ODs=1-3. The following day, 1 mL samples were aliquoted from the overnight cultures and were inoculated with 5 mg/L zillamycin. These samples were cultured for 4 hours at 37C. Following culturing, each strain was spun down and washed with fresh PBS for 3x times. Following the third wash, these samples were extracted in methanol and injected onto the mass spectrometer.

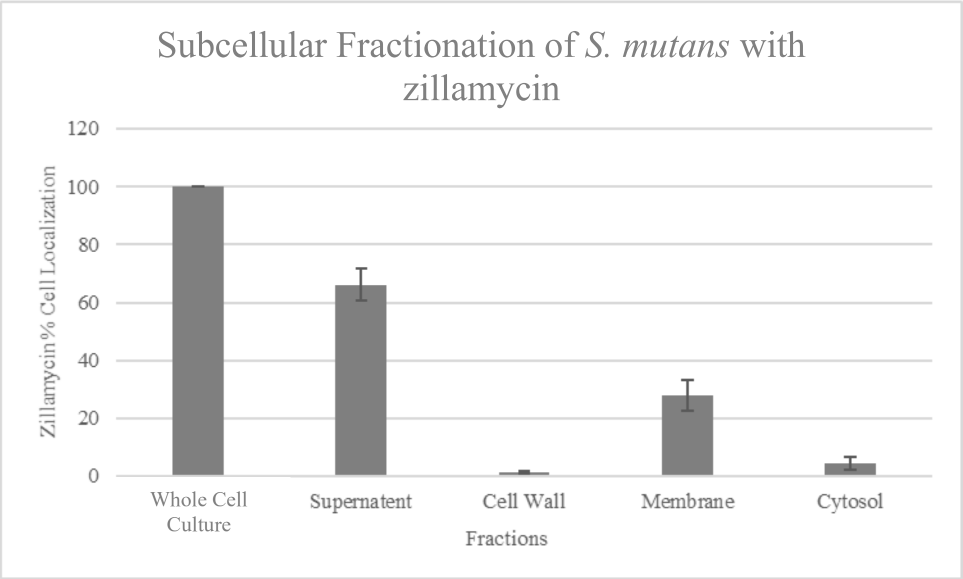

Figure S9: Subcellular fractionation of *Streptococcus mutan*s incubated for 4 hours with zillamycin. Results indicate that the bound compound significantly localizes with the cellular membrane.

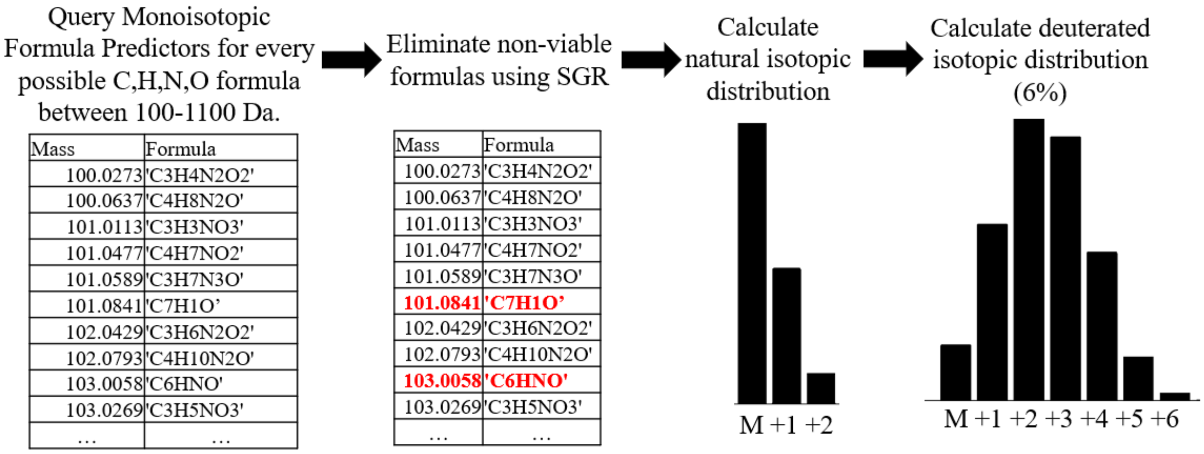

Figure S10: Process for generating a comprehensive mass and formula list with isotopologue abundances. A combinatorial list of the possible formulas containing carbon, hydrogen, nitrogen, and oxygen from masses 100-1100 Dalton was generated. This list was paired down for "realistic" formulas by referencing rules set out in the "Seven Golden Rules" paper. Specifically, the number of each element allowed in a mass range, the following of LEWIS rules, H/C ratios, H/N, and H/O ratios were all modulated to reflect within three standard deviations of existing natural products. From the remaining formulas, both natural and deuterated (6%) isotopic abundances were calculated.

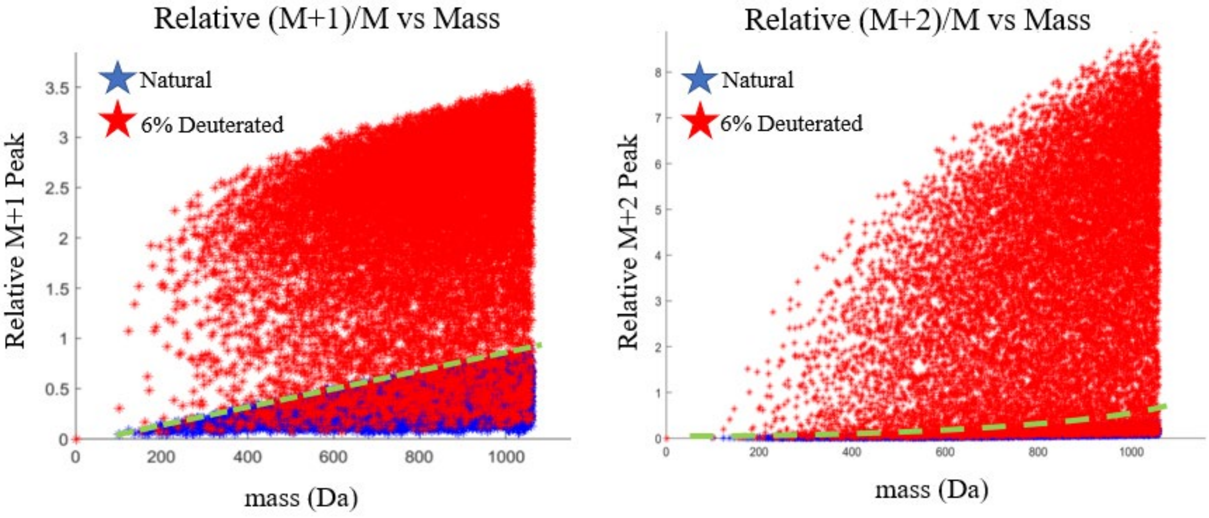

Figure S11: Relative mass difference between a) (M+1)/M and mass or b) (M+2)/M and mass. Plotted in both graphs are both the natural distribution (blue star) and the 6% deuterated distribution (red star) of the isotopologues. The plotted green line shows the maxima of the natural region. Approximately 93.5% of the deuterated formulas fall above the green line within their given mass region for both graphs.

NMR Discussion

Structural Elucidation of zillamycin. Zillamycin had the molecular formula $C_{31}H_{53}N_3$ based on a proton adduct ion at m/z 468.4313 $[M+H]^+$ in its positive ion HRMS spectrum. Analysis of its $^1H$ NMR, and $^{13}C$ NMR spectroscopic data of compound 468 (Table S5) indicated that compound 468 included one guanidine moiety and six isoprene units. Furthermore, the structure of compound 468 was confirmed as a triterpene guanidine through investigating its COSY and HMBC spectra.

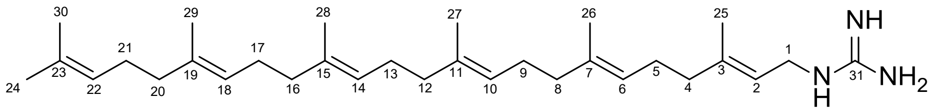

Additional Methods and Materials

Mass spectrometry

Extracted metabolites were run on an Agilent QToF 6545 mass spectrometer with an Agilent C18 analytical column and an Agilent 1240 DAD detector upstream. The instrument is run at 100-1700 Da mass range and the fragmenter is set to 110 V. These files are recorded by Agilent Mass Hunter Workstation Data Acquisition Version B.09.00.

HPLC

Concentrated extracts were purified using an Agilent 1260 HPLC with a Sephadex C18 column and an Agilent 1260 DAD detector upstream.

NMR

A 900 MHz NMR was utilized to solve structures. Approximately 1 mg of compounds was suspended in $CDCl_3$. Structures were solved by Dr. Yongle Du.

Various Resins

For size exclusion chromatography, a packed column of Sephadex LH20 resin with an eluent of methanol. For packed reverse phase chromatography, C18 SP columns were utilized.

$D_2O$ Sorter Code

Below is the "$D_2O$ Sorter" code used to identify deuterated small molecules written in MATLAB:

```
file=xlsread('Insert Excel File Name')
%file=readtable([namesave{po} '.txt'],'Delimiter', '  ');

%Parameters
error1=0.1;%
RTerror=.08; %0.08mins, allowed retention time error/dalton
MinAbund=0; %Minimum allowed abundance
ratio1toM=0.09;
ratio2to1=0.01;

if(individual==true)
%file=file(3:end,[2,3,7]);
%file(:,[1 2])=file(:,[2 1]);
mzdata=[];
mzdata(:,1:3)=file(:,1:3);
mzdata=sortrows(mzdata,1);%Sorts the m/z's into rention time order and matches them with their
according intensity value
elseif(individual==false)
file=table2array((file(4:end,[2,3,27])))
file(:,[1 2 3])=file(:,[2 1 3])
mzdata=[];
mzdata(:,1:3)=cell2mat(file(:,1:3));
mzdata=sortrows(mzdata,1)
end

%For Specific Searching
if(true)
mzlimit=[400,500];
rtlimit=[33,37];
mzdata(mzdata(:,1)>mzlimit(2),:)=[];
mzdata(mzdata(:,1)<mzlimit(1),:)=[];
mzdata(mzdata(:,2)>rtlimit(2),:)=[];
mzdata(mzdata(:,2)<rtlimit(1),:)=[];
end

%MZ Isotope Sort

mzlist=zeros(60,length(mzdata));
abundancelist=zeros(60,length(mzdata));
rtlist=zeros(60,length(mzdata));
```

```
 for sw=1:length(mzdata);%
     mztemp=zeros(45,1);
     abundtemp=zeros(45,1);
     rttemp=zeros(45,1);

     mztemp(1,1)=mzdata(sw,1);
     abundtemp(1,1)=mzdata(sw,3);
     rttemp(1,1)=mzdata(sw,2);

     error=error1; %%gives a ppm error range

        rtdiff=abs(mzdata(sw,2)-mzdata(:,2));
        integerdiff=round(mzdata(:,1)-mzdata(sw,1));
        rtmatchesself =rtdiff<RTerror*integerdiff; %& rtdiff>-.005; %makes the triangle that gives
a larger range with biger integer differences
        mzmatches = abs(mzdata(:,1)-mzdata(sw,1)- abs(1.00627*round(mzdata(:,1)-
mzdata(sw,1))))<error & mzdata(:,1)-mzdata(sw,1)<10;

   %Here we want to ensure that the M has a viable M+1 and M+2 before it gets sorted with our
higher order ones.
        abund1match=((mzdata(:,3).*(integerdiff==1)))>0;
        correctabund1match=abund1match&mzmatches&rtmatchesself;
        abund2match=((mzdata(:,3).*(integerdiff==2)))>0;
        correctabund2match=abund2match&mzmatches&rtmatchesself;

    abundboth=any(correctabund1match)&any(correctabund2match);

       if(abundboth)
       up1=find(correctabund1match&rtmatchesself,1);
       mida1=rtmatchesself(sw+1:up1-1);
       up2=find(correctabund2match&rtmatchesself,1);
       mida2=rtmatchesself(up1+1:up2-1,1);
       midders=any(mida1)&any(mida2);
       else
       midders=0;
       end

       % if()
      %  mids1=mzdata(mzdata(:,1)<up1(1)&mzdata(:,1)>mzdata(sw,1),:)-mzdata(sw,1);  %These
lines produce comparable values between 0-1 for us to compare fragments
      % mids2=mzdata(mzdata(:,1)<up2(1)&mzdata(:,1)>up1(1),:)-up1(1);
      % mids3=mzdata(mzdata(:,1)<up3(1)&mzdata(:,1)>up2(1),:)-up2(1);
      % end
      % end
```

```
        all=rtmatchesself&mzmatches;

      if(any(mzdata(all,2)~=mzdata(sw,2)))
        tooperfect=false;
      else
        tooperfect=true;
      end

        if(any(all)&abundboth&~midders&~tooperfect) %accounts for xcms fuck-ups and averages
in rt/mz based on abundance and adds abundances

             position=round(mzdata(all,1)-mzdata(sw,1));
             mztemp(position+1,1)=mzdata(all,1);
             rttemp(position+1,1)=mzdata(all,2);
             abundtemp(position+1,1)=mzdata(all,3);
             mzdata(all,:)=0;
        end

       mzlist(1:length(mztemp),sw)=mztemp;
       abundancelist(1:length(abundtemp),sw)=abundtemp;
       rtlist(1:length(rttemp),sw)=rttemp;

 end

     %This portion removes all of the zeros from the code

%gets rid of rows with all zeros
abundancelist(~any(mzlist,2),:)=[];
rtlist(~any(mzlist,2),:)=[];
mzlist(~any(mzlist,2),:)=[];

abundancelist(:,~any(mzlist,1))=[];  %gets rid of columns with any zeros
rtlist(:,~any(mzlist,1))=[];
mzlist(:,~any(mzlist,1))=[];

 % with all the values that made it past retention time
%correction stuff
AAA=[];
if(size(mzlist,1)>1)
AAA(:,1)=mzlist(1,:)';
```

```
AAA(:,2)=sum(mzlist~=0,1)';
AAA(:,3)=rtlist(1,:)'; %averages their two retention times for display
AAA(:,4)=abundancelist(1,:)'; %averages abundances for display
AAA(:,5)=(abundancelist(2,:))';
AAA(:,6)=(abundancelist(3,:))';
if(size(abundancelist,1)>3)
AAA(:,7)=(abundancelist(4,:))';
end
%AAA(:,8)=(abundancelist(5,:))';
AAA(:,8)=AAA(:,4)./AAA(:,4);
AAA(:,9)=AAA(:,5)./AAA(:,4);
AAA(:,10)=AAA(:,6)./AAA(:,4);
AAA(:,11)=AAA(:,7)./AAA(:,4);

AAA(:,15)=sum(abundancelist(1:end,:))';
AAA(:,16)=0.00078254.*AAA(:,1)+1*0.0476934;
AAA(:,17)=0.000000287705706*(AAA(:,1).^2)+0.000039169586817*AAA(:,1)+1*0.00172339
7268419;
AAA(:,18)=0.00115025.*AAA(:,1)+1*2.3113929;
AAA(:,19)=-
0.000001637010966*(AAA(:,1).^2)+0.008100842649*AAA(:,1)+1*2.087517543739501;

AAA(AAA(:,2)<2,:)=[]; %removes all samples outside of abundance range

else
AAA=[]
end

% Applies Formula Removals (using my max molecular formula predictor)
  min=(AAA(:,5)./AAA(:,4))<AAA(:,16)|(AAA(:,6)./AAA(:,4))<AAA(:,17);

%min=(AAA(:,5)./AAA(:,4))<realistic*peaktempsmin(:,2)|(AAA(:,6)./AAA(:,4))<realistic*peak
tempsmin(:,3)|(AAA(:,7)./AAA(:,4))<realistic*peaktempsmin(:,4);
  max=(AAA(:,5)./AAA(:,4))>AAA(:,18)|(AAA(:,6)./AAA(:,4))>AAA(:,19);

%max=AAA(:,6)>1.1*peaktempsmax(:,2)|AAA(:,7)>1.1*peaktempsmax(:,3)|AAA(:,8)>1.1*pea
ktempsmax(:,4)|AAA(:,9)>1.1*peaktempsmax(:,5)

eval(['AAA_' namesave{po} '=AAA']);
eval(['AA_fake_' namesave{po} '=AA_fake']);
eval(['AA_real_' namesave{po} '=AA_real']);
eval([namesave{po} '_results={AA_fake_' namesave{po} ', AA_real_' namesave{po} ', AAA_'
namesave{po} ', peaktempsmin}' ])
```

```
eval(['save ' namesave{po} '_results ' namesave{po} '_results'])

namesave{po}

end
%% Table
for poq=1
%Makes an actual table
r={'M','# ++', '# 1-3', 'RT','Abundance M','Abund M+1/M','Abund M+2/M+1','RT slope','R^2'};
f=figure('position',[600 800 300 500],'units','normalized','outerposition',[0 0 1 1]);
uitable(f,'data',AAA,'columnname',r,'position',[0 0 700 600],'ColumnWidth', 'auto')

toc;
end
```
